## Supplemental Fig 1 for "In vitro evidence against productive SARS-CoV-2 infection of human testicular cells: Bystander effects of infection mediate testicular injury"

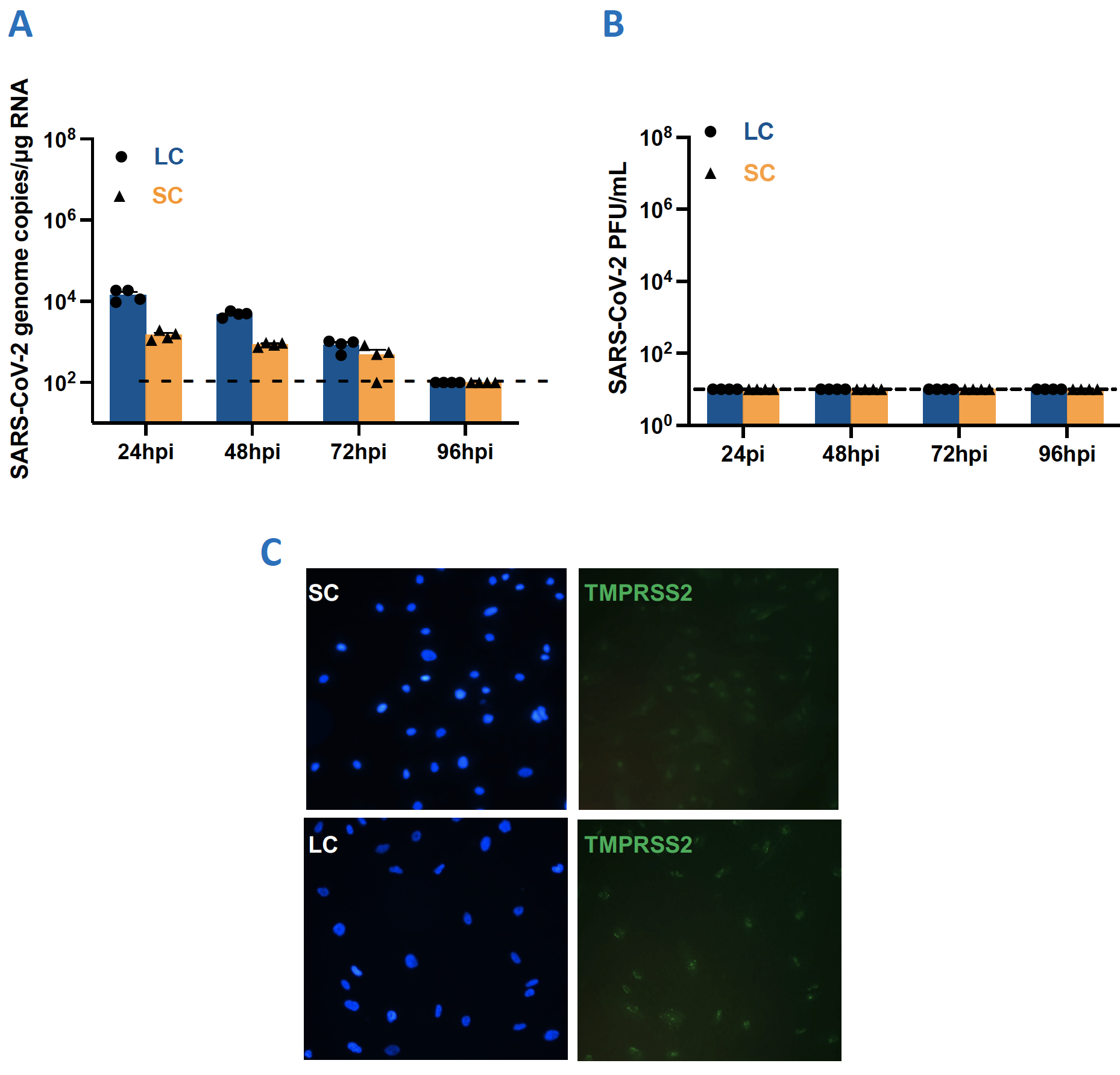

**Supplement Figure 1:** Primary SC, LC, STC, and HTO were infected with SARS-CoV-2 at MOI 10 and **(A)** intracellular virus levels were determined using qRT-PCR and **(B)** virus titers in the supernatant were measured using plaque assay **(C)** Representative TMPRSS2 (green) and DAPI (blue) staining of primary Sertoli (top) and Leydig (bottom) cells.
